# Dual-Drug Loaded Biomimetic Chitosan-Collagen Hybrid Nanocomposite Scaffolds for Ameliorating Potential Tissue Regeneration in Diabetic Wounds

**DOI:** 10.1101/2022.02.16.480700

**Authors:** Vyshnavi Tallapaneni, Divya Pamu, Lavanya Mude

## Abstract

Diabetes Mellitus (DM) is one of the most concerning conditions, and its chronic complications are nearly synonymous with inflammation, oxidative stress, and infections. In the acute inflammatory phase of diabetic wound healing (DWH), reducing excessive reactive oxygen species (ROS) and inflammatory response of the wound is a necessary treatment. The current work used a mix of emulsification and lyophilization approaches to investigate the effects of resveratrol microparticles (RES-GMS) loaded chitosan-collagen (CS-CLG) scaffold with doxycycline (DOX) on DWH. Resveratrol (RES) is a powerful antioxidant that promotes cell proliferation in the dermis by improving fibroblast function and enhancing CLG production. DOX can potentially shift the balance away from the chronic wound’s pro-inflammatory, proteolytic status toward an environment that promotes vascular ingrowth and, eventually, epithelial development. Cross-linked scaffolds had optimal porosity, reduced matrix degradation, and prolonged drug release when compared to non-cross-linked scaffolds, according to the results of composite scaffold characterization. Cell proliferation assay employing mouse fibroblasts was used to study the kinetics and bioactivity of growth factors produced from the scaffold. The RES-DOX-CS-CLG scaffold was biocompatible and promoted cell development compared to the control and CS-CLG scaffolds in *in vitro* experiments. DOX-loaded CS-CLG scaffold loaded with R-GMS delivers a prolonged release of RES, according to *in vitro* tests.

## Introduction

Diabetes mellitus (DM) is a class of metabolic diseases defined by hyperglycemia as a consequence of insulin resistance or insufficiency. Long-duration high blood glucose levels can lead to organ dysfunction and failure. Giving the World Health Organization (WHO), 422 million people have diabetes, with 200 million individuals suffering from short- and long-term consequences worldwide (1, 2). Non-healing wounds, including DWs, are one of the most prevalent significant consequences of diabetes. These wounds can cause lower quality of life, serious infections, and amputations (3, 4). DWs that do not heal are the most common reason for hospitalization, and the overall cost of DWs to the healthcare system is more than US$25 billion annually. Multiple variables, including aberrant inflammatory retention and a diminished granulation tissue condition, cause wound healing to be delayed in people with diabetes. These factors primarily result from reduced glucose metabolic rate and neurovascular difficulties (5–7).

DWs are hard to repair and tend to become chronic or stalled. The wound bed microenvironment of a chronic or halted wound differs significantly from that of an acute or healing wound, with the high inflammatory intermediaries and proteases levels (8, 9). These wounds will still be in the inflammatory phase, marked by a constant influx of neutrophils and the production of cytotoxic enzymes, reactive oxygen species (ROS), and inflammatory mediators that cause tissue damage. In the presence of ROS, defense against invading pathogens occurs, and low ROS levels are essential for intracellular signaling (10). Excessive ROS levels, such as those found in inflammatory diabetic wounds, cause leukocyte activation and endothelial cell destruction. Antioxidant enzymes such as catalase, superoxide dismutase, and glutathione peroxidase compensate for ROS generation from mitochondria in cells without oxidative stress. When the oxidative equilibrium is disrupted, ROS can cause abnormalities in normal cellular activities as well as the breakdown of cells (11, 12). As a result, neutralizing free radicals using an antioxidant and reducing/terminating chronic inflammation may be a powerful technique for accelerating wound healing in DWs. RES is a powerful antioxidant that promotes cell proliferation in the dermis by improving fibroblast function and enhancing CLG production. As a result, using RES to improve the slow healing of diabetic wounds could be a viable option (13, 14).

By coordinating epithelialization and angiogenesis, the ECM plays a vital role in wound healing. Beyond contributing to tissue structural integrity and strength, the matrix provides the basis for vascular endothelial cell ingrowth and a sufficient substrate for keratinocyte migration and adhesion (15, 16). To date, the active dressings that have evolved are either focused on treating the infection by delivering antimicrobials, or they aim to prevent infection by administering antimicrobials. Wound substitutes now on the market have many drawbacks, including wound contraction, scar development, and poor tissue integration (17). The three-dimensional scaffolds can be utilized to cover wounds, provide a physical barrier against infection as a wound dressing, and sustain dermal fibroblasts and overlying keratinocytes for skin tissue engineering (18). An exemplary tissue scaffold should have the proper physical and mechanical capabilities and the right surface chemistry and nano and microstructures to facilitate cellular attachment, proliferation, and differentiation (19).

Due to their ease of manufacture and possible applications, CLG-based products have become increasingly popular in tissue engineering, particularly diabetic wound healing. They come in different forms, including films, gels, fibres, and sponges. Porous 3D CLG scaffolds have received more attention among these many types because they facilitate cell-biomaterial interactions, cell adhesion, and ECM deposition (20, 21). Another significant disadvantage of CLG as a scaffold is its bio-instability. Chemical cross-linking or combined hybridization with synthetic polymers or natural polysaccharides are considered effective methodologies for manufacturing superior CLG-based scaffolds to minimize the possibility of CLG degradation and improve its low mechanical strength. Natural polymers are preferred over synthetic polymers because of their outstanding biodegradability and biocompatibility (22) (23). Numerous co- and synthetic polymers have been employed in conjunction with CLG. As a result, in this work, CS was combined with CLG and then cross-linked to increase its physical stability while simultaneously providing a moist wound environment (24).

DOX can potentially shift the balance away from the chronic wound’s pro-inflammatory, proteolytic status toward an environment that promotes vascular ingrowth and, eventually, epithelial development. A research paper documenting the wound healing capabilities of RES-DOX using CS-CLG, has yet to be found to the best of our knowledge (25). CLG and CS-based products have grown in popularity in tissue engineering, particularly in diabetic wound healing, due to their ease of manufacture and potential wound healing applications. In light of the above research, a combination of topical anti-inflammatory, antioxidant and antibiotic agents, i.e., RES and DOX, can be loaded into the CS-CLG scaffold in this current study, which may effectively address the complex primary pathophysiology of DWs, resulting in synergistic wound repair. These RES-GMS-CS-CLG scaffolds help diabetic wounds heal faster by reducing oxidative stress, inflammation and infection while also promoting cell proliferation and tissue regeneration (26–28).

## MATERIALS AND METHODS

### Materials

Doxycycline Hyclate, Collagen (Porcine type I), gelatin, and Chitosan (MW 100-300 kDa) were purchased from MP Biomedicals (India) Pvt Ltd, Mumbai, 2-morpholinoethane sulfonic acid (MES), Sodium tripolyphosphate (TPP), 1-ethyl-(3-3-dimethyl aminopropyl) carbodiimide hydrochloride (EDC, MW=191.7), N-hydroxysuccinimide (NHS), Collagenase, Triethylamine, Acetonitrile and Methanol of HPLC grade were procured from Sigma Aldrich, USA. Trypsin, Dulbecco’s modified essential medium (DMEM), and Foetal bovine serum (FBS) were purchased from Himedia, India.

### Preformulation studies

#### Compatibility studies

The DSC Q200 was used to investigate the drug compatibility of the chosen medicines (RES and DOX) (TA Instruments, USA). The samples were enclosed in aluminum pans and heated (100C per minute, 30–3000C temperature) under a 40 mL/min nitrogen flow with an empty pan as reference. Thermograms for RES, DOX, and their physical combination were obtained (15).

#### Preparation of RES loaded gelatin microspheres (RES-GMS)

To make RES-GMS, the emulsification-linkage process was applied. RES was solubilized in a pre-swelled gelatin solution, then poured into liquid paraffin (55 1°C) to form a w/o emulsion. Then followed by the addition of glutaraldehyde as cross-linker of 25 percent to the system to cross-link the gelatin droplets for 30 minutes. Drying was done by vacuum filtration and triple washed with petroleum ether and isopropyl alcohol (29).

### Characterization of RES-GMS microspheres

#### Encapsulation efficiency

The EE of RES within the drug-loaded GMS was evaluated by measuring the absorption of the clear supernatant using a UV-spectrophotometer after the suspensions of the drug-loaded microspheres were centrifuged at 17,000 rpm for 40 minutes (REMI R-8C, India). Testing the supernatant of blank microspheres yielded the relevant calibration curves. For each sample, tests were carried out in triplicate. A UV–vis spectrophotometer was used to determine the absorbance value of RES at a wavelength of 304 nm (30).

The amount of RES encapsulated in microspheres is expressed as EE% and calculated as follows.

EE (%) = (Mass of the drug in nanoparticle) / (Mass of the drug used in the formulation) X 100

#### Determination of particle size (PS), zeta potential (ZP), and polydispersity index

The particle size analyzer (Litesizer 500, UK) was used to determine the average size and size distribution of the drug-loaded NPs. To verify that the particles were well disseminated and the dispersion was homogenous, samples measurements were produced by diluting the microspheres suspension with deionized water and sonicating for 30 seconds before testing.

#### Texture analysis

SEM is used to verify the RES-GMS morphology (size and shape). Microspheres are re-suspended in distilled water before being placed on a silicon grid and allowed to dry at ambient temperature. Before SEM examination, the suspension was vacuum coated with gold for 3 minutes. The surface properties of the samples were examined using a Carl Zeiss SEM (Germany) with a 15-keV pulse and various resolutions (31).

#### Formulation of CS-CLG scaffolds impregnated with RES-GMS and DOX (RES-GMS-CS-CLG scaffolds)

Scaffolds were prepared using the freeze-drying process. CLG solution (4 wt%) is generated by dissolving CLG powder in 0.5 M acetic acid, while CS solution (4 wt%) is made by dissolving CS powder in distilled water. The CS- CLG blends are next made; first, the pH of acidic CLG is corrected to 7 at 4°C by adding 2 M NaOH, then the CS solution is added dropwise to the CLG solution, and the final pH is adjusted to 6-7. Finally, after 2 hours of continuous stirring, a precise homogeneous blend is formed, centrifuged for 15 minutes at 4000 rpm to remove entrapped air bubbles. The resultant mixture is then poured into molds to create hydrogels. After being rinsed with deionized water, the hydrogels are put in polystyrene culture flasks and lyophilized at −800C for 72 hours, resulting in porous CLG matrices. CS- CLG composite scaffolds with a 50/50 (w/w) CS-CLG ratio were made and then expressed as CS- CLG scaffolds (Placebo scaffolds). The scaffold discs are chemically cross-linked with EDC/NHS (EDC-cross-linked) for 24 hours at 4°C in MES buffer solution (pH 5.5, 0.05 M) (19). To make drug-loaded scaffolds, RES-GMS and DOX are mixed in a CS solution with magnetic stirring, then dropped into a CLG solution to achieve a final concentration of 1% (w/v) RES and 1% (w/v) DOX. The drug-loaded CS- CLG composite mixture is then agitated for an overnight period to achieve uniform mixing of the RES-GMS and DOX, before being placed into a culture flask, frozen, and lyophilized. For further testing, the freeze-dried scaffolds are kept in a desiccator. RES-GMS and DOX-loaded CS- CLG scaffolds will be referred to as RES-GMS-CS-CLG scaffolds from now on (32).

### Characterization of RES-GMS-CS-CLG scaffolds

#### Texture morphology

A scanning electron microscope was used to examine the surface architecture of fibrin-based scaffolds (FlexSEM 1000, Hitachi, Tokyo, Japan). Scaffold samples were totally dried after 30 minutes of incubation at 37°C to accomplish fibrin polymerization, hence no extra procedures were used before SEM examination. With a 5 kV acceleration voltage, SEM microphotographs were obtained at 5000 to 20000X magnifications. An open-source image processing application was used to examine the photos (ImageJ). Six random measurements were taken for each picture to establish the mean diameter of the fibrin nanofibers. Pore areas were also computed in SEM images using a subjective approximation of surface pores (30 measurements calculated areas) (31).

#### Tensile strength measurement

Mechanical characterization was performed on scaffolds (10 × 20 mm2) using a tensile testing equipment (Tinius Olsen, H5KS Model) with a 5 KN load creating a cell. Experiments on tensile strength were carried out at room temperature at a stretch rate of 1 mm/min. Young’s modulus and ultimate tensile strengths were calculated using the obtained force and extension values. All data are mean ± standard deviation (n=3) (27).

#### Water uptake

W_dry_ is the weight of the cross-linked and non-cross-linked RES-GMS-CS-CLG scaffold discs [8 mm diameter and 1.5 mm thickness]. After that, the scaffold samples are soaked in a closed tube containing 5mL SWF at a pH of 7.4 and kept at 37°C. After 72 hours, the swelled samples are withdrawn from the tube, and the excess liquid is blotted away using filter paper before being weighed as W_wet_ (33).

The following formula is used to compute the water uptake by swelling ratio (S):

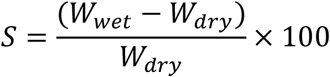

#### Matrix degradation studies

Collagenase degradation is used to examine the in vitro biodegradation of the EDC-cross-linked RES-GMS-CS-CLG scaffold; non-cross-linked scaffolds are also tested for comparison. Analysis of hydroxyproline content provides a measure of the amount of CLG degradation in the treated scaffolds. “The percentage ratio of the hydroxyproline released from the scaffolds at different intervals to the destroyed one with known composition and known weight,” according to the definition of degree of biodegradation. The mean ± standard deviation (n = 3) were used to represent the data (34).

#### *In vitro* drug release studies

*In vitro*, RES is released from GMS using the dialysis bag membrane method. In a dialysis tube, RES-GMS (100 mg) is suspended in 2 mL of phosphate buffer saline (pH 7.4) and maintained at 37°C (MW CO 13 KDa, HiMedia, India). In addition, it was immersed in 200 mL of SWF (pH 7.4, 37°C, constant stirring). By dispersing the composite scaffold (3 cm2 [1.5X2cm]) in 20 mL of SWF, RES, and DOX are released from the RES-DOX-CS-CLG scaffold (pH 7.4 at 37°C). At regular intervals, the supernatant is pipetted out and replaced with equivalent amounts of new phosphate buffer solution. A UV-Visible Spectrophotometer set to 304 nm for RES in ethanol standard curve and 275 nm are used to quantify the quantity of medication released (35).

#### Kinetics of drug release *In vitro*

The release data is fitted to models reflecting zero order, first order, and Higuchi’s square root of time kinetics to analyze the drug release kinetics (36).

#### *In vitro* antibacterial studies

The *in vitro* antibacterial properties of RES-DOX-CS-CLG scaffolds and placebo scaffolds were investigated using the disc diffusion technique. The microorganisms utilized were as follows: Gram-negative Escherichia coli (E. coli) and Pseudomonas aeruginosa (P. aeruginosa), Gram-positive Staphylococcus aureus (S. aureus), and Methicillin-resistant Staphylococcus aureus (MRSA). In a nutshell, a nutrient agar plate with evenly spread bacterial suspension had the samples placed upon its surface & subjected to overnight incubation (37°C) followed by zones of inhibition measurement. The diameter of inhibition zones (in centimetres) measured is shown in the **Table 3**. The clear zone diameter of the bacterial inhibition zone was correlated to antibiotic activity (DOX and Drug loaded formulation) for gram-positive and Gram-negative bacteria, respectively (37).

#### *In vitro* cytotoxicity studies on fibroblast-3T3 cells

To assess the viability of the RES-DOX-Cs-CLG scaffolds, the Sulforhodamine B (SRB) test was utilised. Scaffolds of standard dimensions (5 mm 5 mm 3 mm) were sterilised in 75% ethanol for 30 minutes, then rinsed five times in sterile water for 5 minutes before being placed in 96-well plates with 2 ml of DMEM in each well. The BALB/3T3 cells were then planted in the 96-well plate, either with scaffolds or without. The test was also conducted over 72 hours with a media control to provide for a more accurate comparison. The control and tested scaffolds were photographed using a light microscope (Almicro, Motic®, China). Three times the experiment was conducted (38).

#### *In vitro* scratch assay

The scratch experiment was carried out using Balb/3T3 fibroblast cell lines. To simulate the diabetic situation, the cells were grown in DMEM with 10% PBS and 50 mM glucose. The cells were then planted onto the tissue culture plates as a monolayer at 70-80 percent confluence. Using a 1 mL pipette tip, a scratch was formed on the plate. Various test samples were produced in new medium, put to the plates, and incubated for 24 hours. Cells were then washed and treated using a 4 percent formaldehyde solution (39).

## RESULTS AND DISCUSSION

### Compatibility study using DSC

Compatibility study using DSC of RES has shown the melting point at 262°C and DOX at 184.18°C. The physical mixture has not demonstrated the shift of meting point peaks and found within both drugs’ melting point range **(Fig.1.)**.

**Fig.1.**
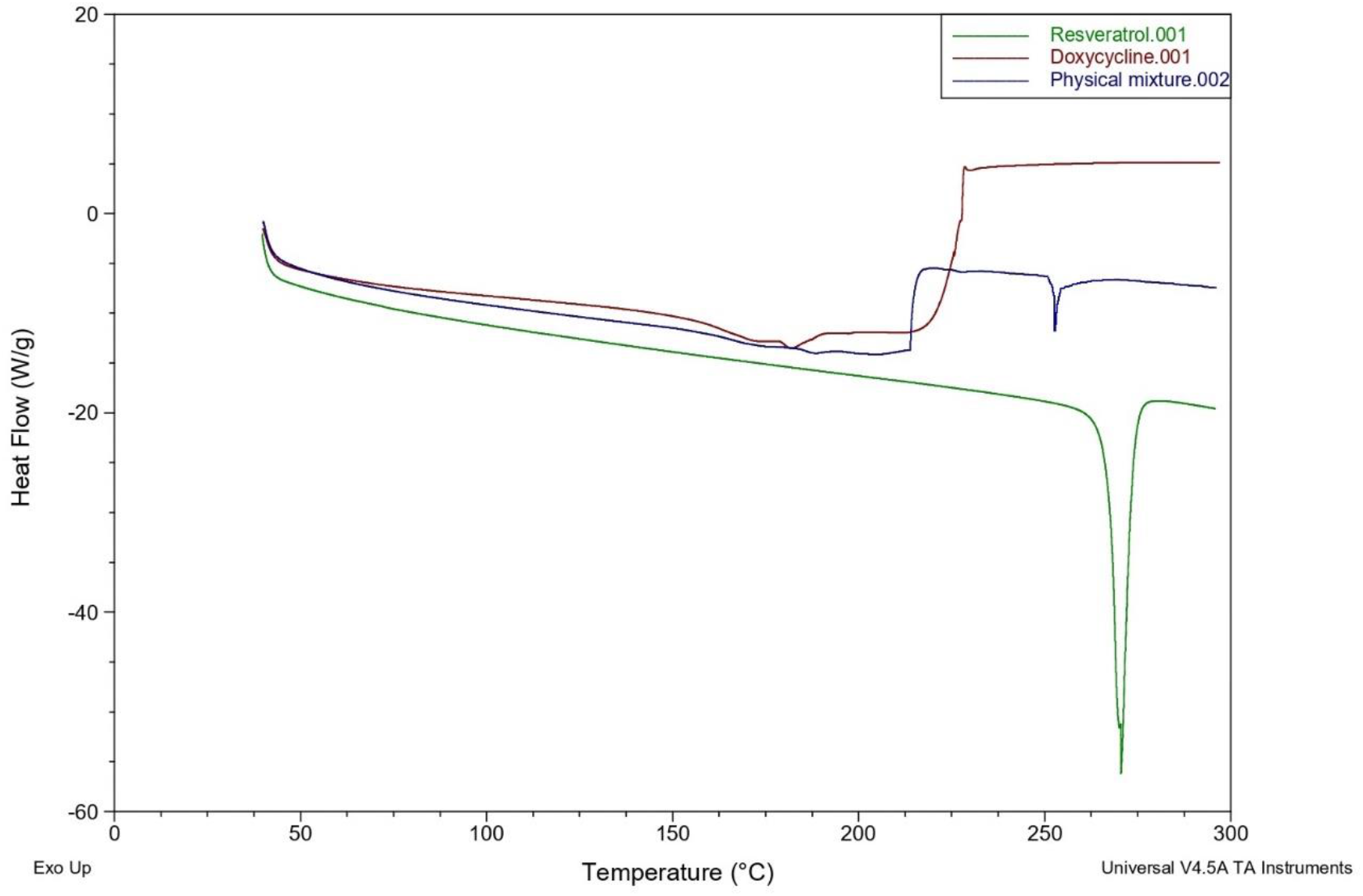
DSC thermograms of Resveratrol, chitosan and physical mixture

### Characterization of R-GMS

RES-GMS of 10mg were carefully weighed and transported to a beaker containing 20 ml of 1 M sodium hydroxide solution, where they were agitated for 12 hours to thoroughly dissolve the microspheres. A UV-Visible spectrophotometer set to 304 nm was used to check for drug content in the produced solution (Shimadzu, Japan). The quantity of RES encapsulated in microspheres was determined and reported as encapsulation efficiency.

The concentration of gelatin and GA solution substantially impacts the EE of microspheres. According to the findings, raising the polymer content from 10% to 20% w/v resulted in a 76-80% increase in EE value. Increasing GA concentration had a favorable impact on EE values as well. Because of the high concentration of GA solution, a stiff network developed, preventing drug molecules from leaking throughout the production process.

The prepared microspheres had an average particle size of 2 um and polydispersity of 0.2 to 2.9. The R-GMS have been found to be spherical in shape with uniform size, homogeneity and micron size **(Fig.2.)**.

**Fig.2.**
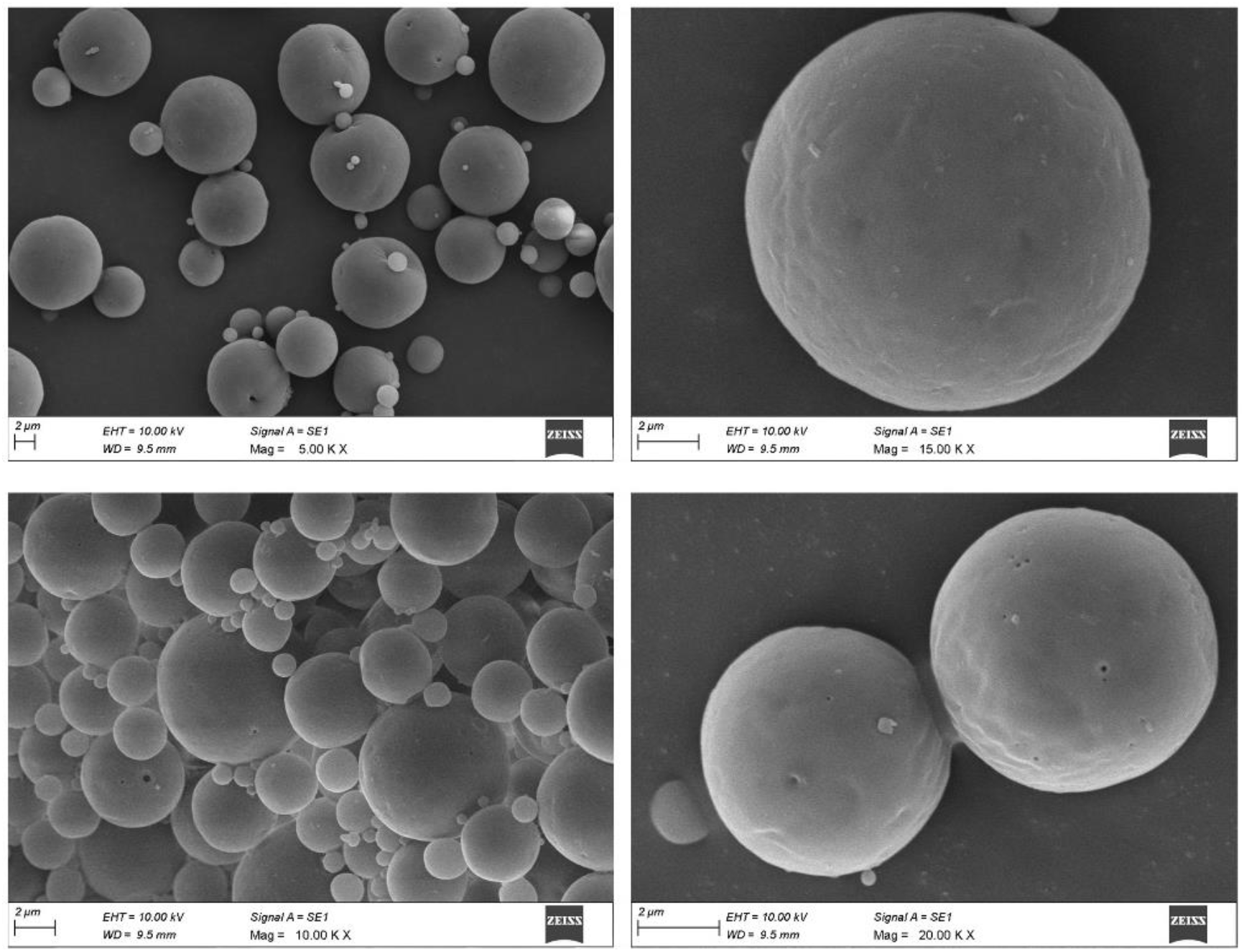
Cross sectional morphologies of RES-GMS using SEM

### Scaffold Characterization

**Fig.3**. shows a lateral slice of RES-DOX-Cs-CLG scaffolds, each of which had a layered morphology due to the mold’s wall structure. To support cell migration and nutrient transfer, the scaffolds should have suitable structural properties such as ideal porosity and pore size, as well as pore interconnectivity. The interconnected macro-scaled channels observed in SEM images **(Fig. 3)** to provide a conduit for the delivery of vital nutrients and metabolic waste into the centre sections of the drug-loaded scaffolds. Freeze drying has the advantage of allowing pore size control in tissue engineering (31). The EDC-crosslinking greatly improved the mechanical characteristics of RES-DOX-Cs-CLG scaffolds compared to CLG and CS-CLG scaffolds suggesting their better *in vivo* stability **(Table 1 & Fig.4.)**.

**Fig.3.**
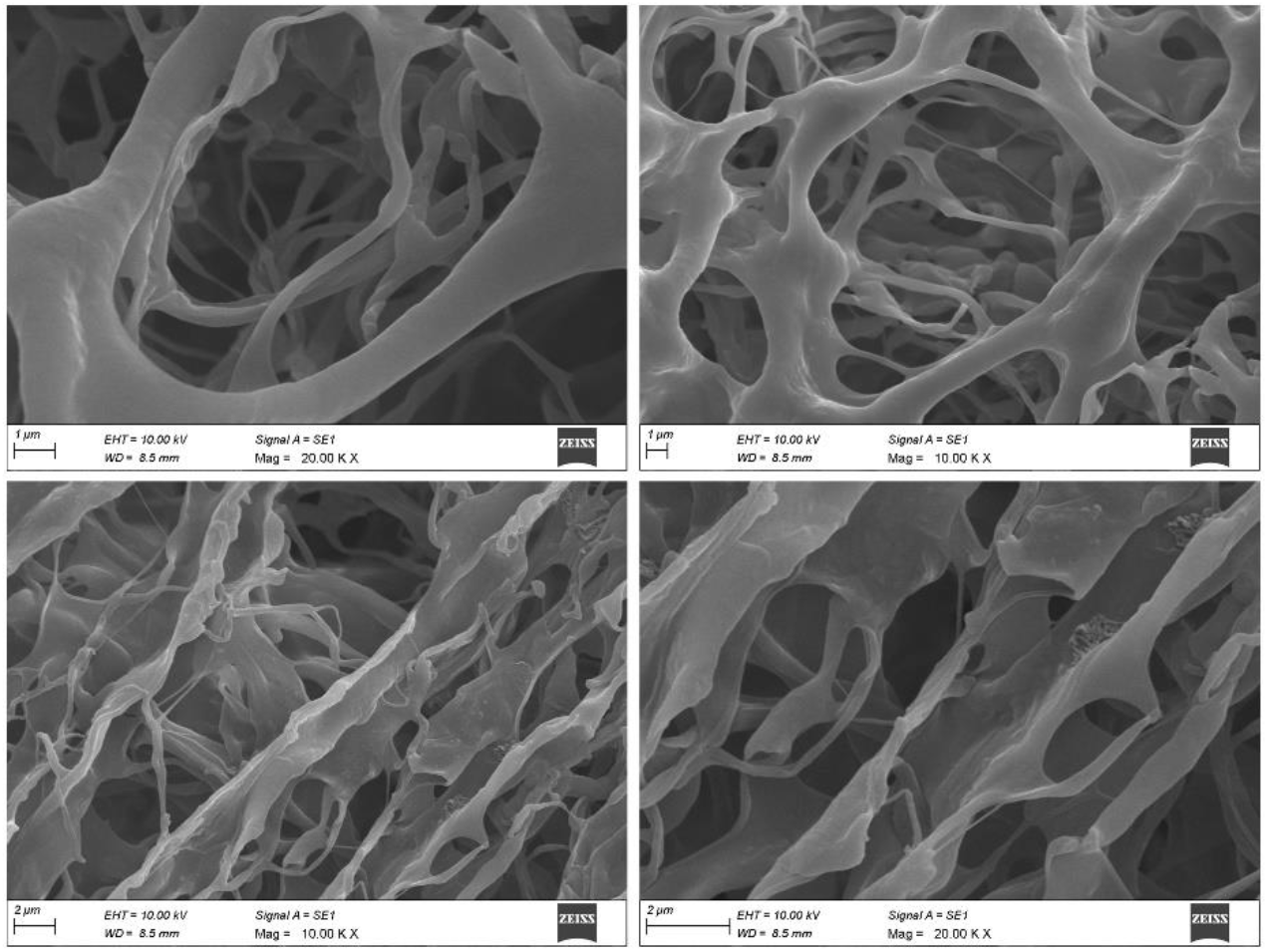
Cross sectional morphologies of scaffolds using SEM

**Table 1.** Comparison of tensile strengths and Young’s modulus among various scaffolds

| Scaffold | Tensile Strength (MPa) | Young's Module (MPa) |
| --- | --- | --- |
| RES-GMS-CS-CLG scaffold | 1.95±0.53 | 13.67±1.2 |
| CS-CLG Scaffold | 0.75±0.99 | 8.79±0.87 |
| CLG Scaffold | 0.25±0.68 | 1.65±0.21 |

**Fig.4.**
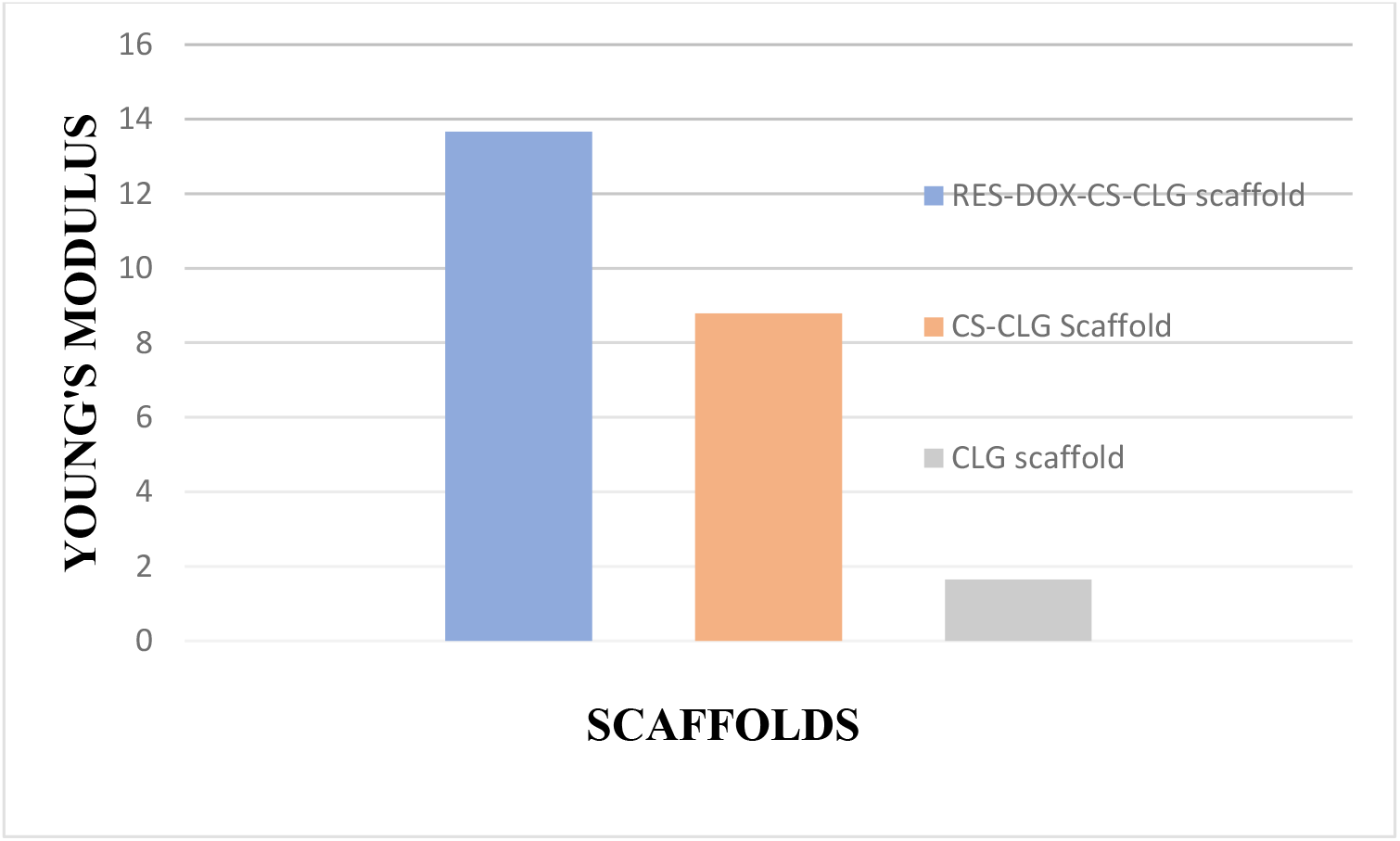
Comparison of Young’s modulus among various scaffolds

The swelling ratios of cross-linked and non-cross-linked scaffolds were determined to be 750 percent and 890 percent, respectively. The swelling ratio was somewhat reduced (140 percent) after cross-linking of RES-GMS-CS-CLG scaffold, most likely owing to the establishment of molecular solid bonds by EDC-crosslinking. Nonetheless, more water absorption may result in a loss of physical integrity and, as a result, a loss of stability throughout the cell culture process and in vivo application **(Fig.5.)**.

**Fig.5.**
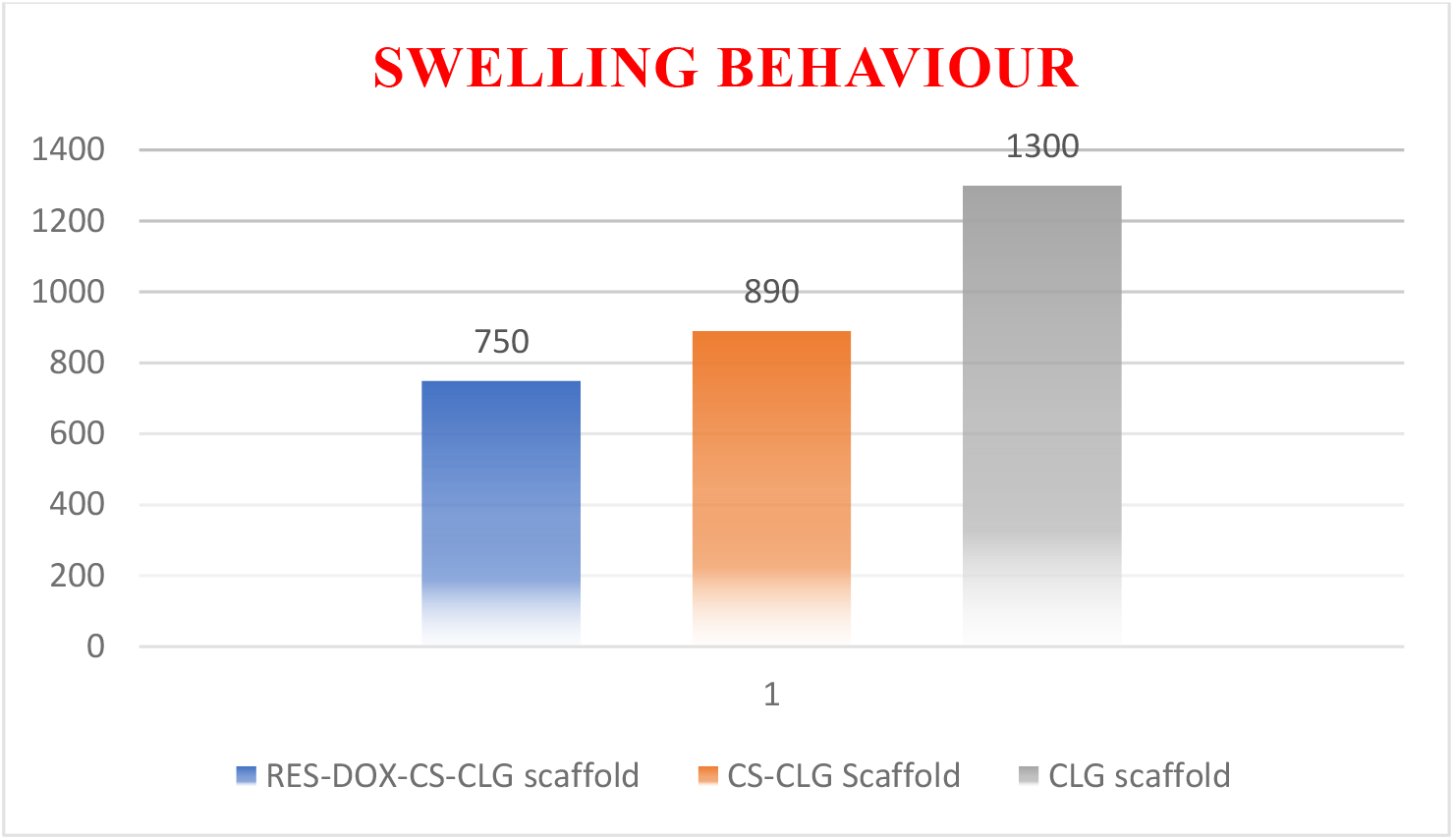
Water uptake of scaffolds before and after crosslinking treatment in pH 7.4 PBS at 37°C

The degradation rate and degree of degradation were significantly reduced when the CS-CLG scaffold was cross-linked to EDC (RES-DOX-Cs-CLG scaffold), implying greater resistance to enzymatic degradation. The cross-linked RES-GMS-CS-CLG scaffold demonstrated the least degradation degree of 22.63 percent after 7 days of incubation. The non-cross-linked CS- CLG and CLG scaffolds showed 92.51 percent and 99.3 percent degradation, respectively **(Fig.6)**.

**Fig.6.**
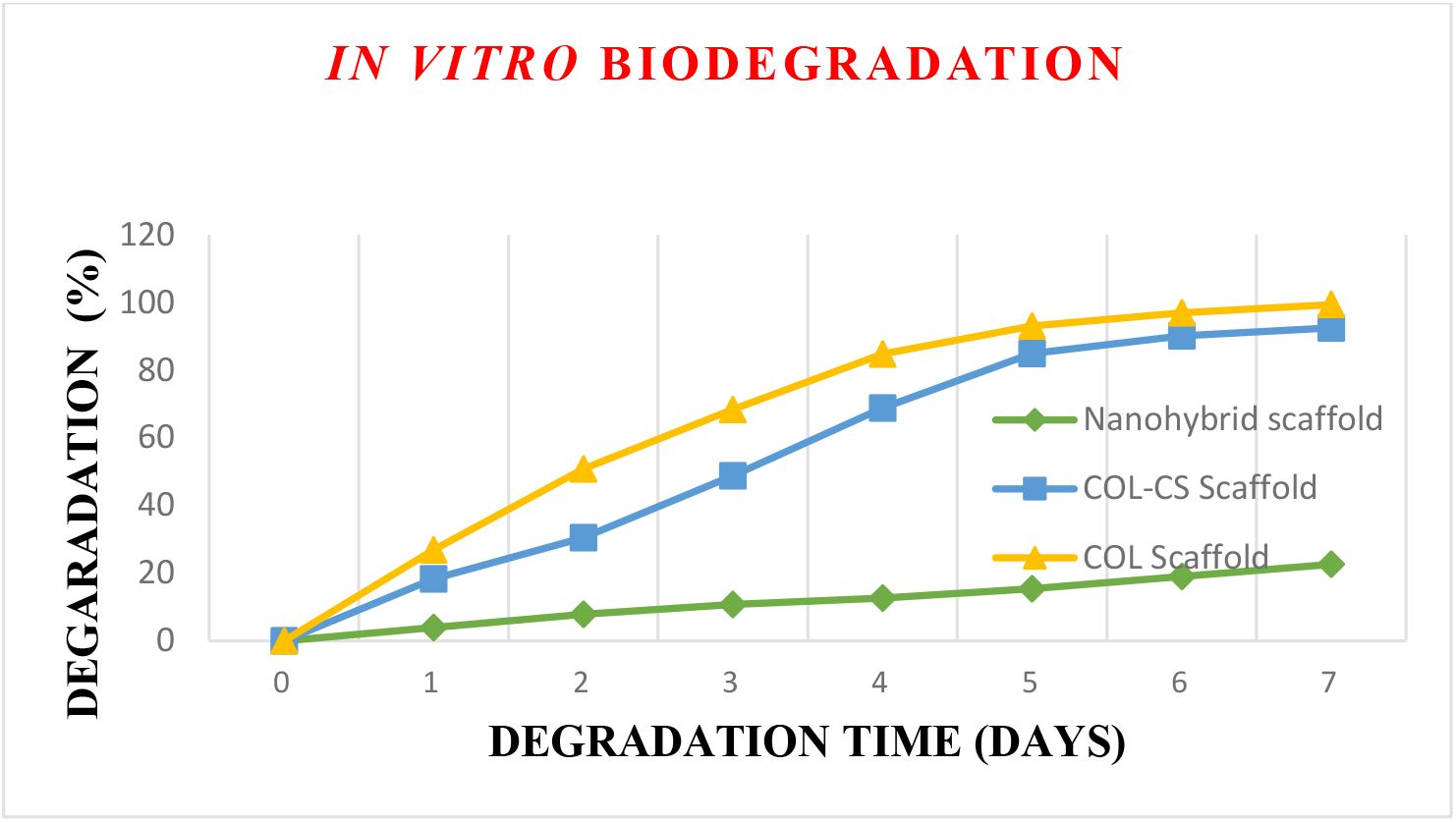
Enzymatic degradation curves of different scaffolds in pH 7.4 PBS containing 265 Umg^−1^ collagenases.

The release of RES and DOX from the RES-DOX-CS-CLG scaffold was achieved by dispersing the composite scaffold (3 cm2 [1.5X2cm]) in 20 mL of SWF (pH 7.4 at 37oC). At regular intervals, the supernatant was pipetted out and replaced with comparable amounts of new phosphate buffer solution. UV analysis at 304 and 274 nm was used to determine the extent of drug release using a standard curve of RES in ethanol and DOX in PH 7.4.

The drug release from RES-GMS established a two-phase tendency marked by fast release for 24h that was accompanied by a sustained release phase. 34.99 % of drug was released from R-GMS whereas 21.57 % of drug was only released from RES-CS-CLG scaffold. Evidently scaffold sustained drug release initially. At 72 hrs of study 55% of drug only had released from the scaffold whereas 80% had released from microspheres meeting the objective stating that it is beneficial only if the delivery system (scaffold) sustains the release of drug during the whole period of treatment. 92 % of drug release from the scaffold at 300h will guarantee effective release of drug over 14 days and above which will minimize the frequency of applications of scaffold and thereby contributes to an effective DWH **(Fig.7)**.

**Fig.7.**
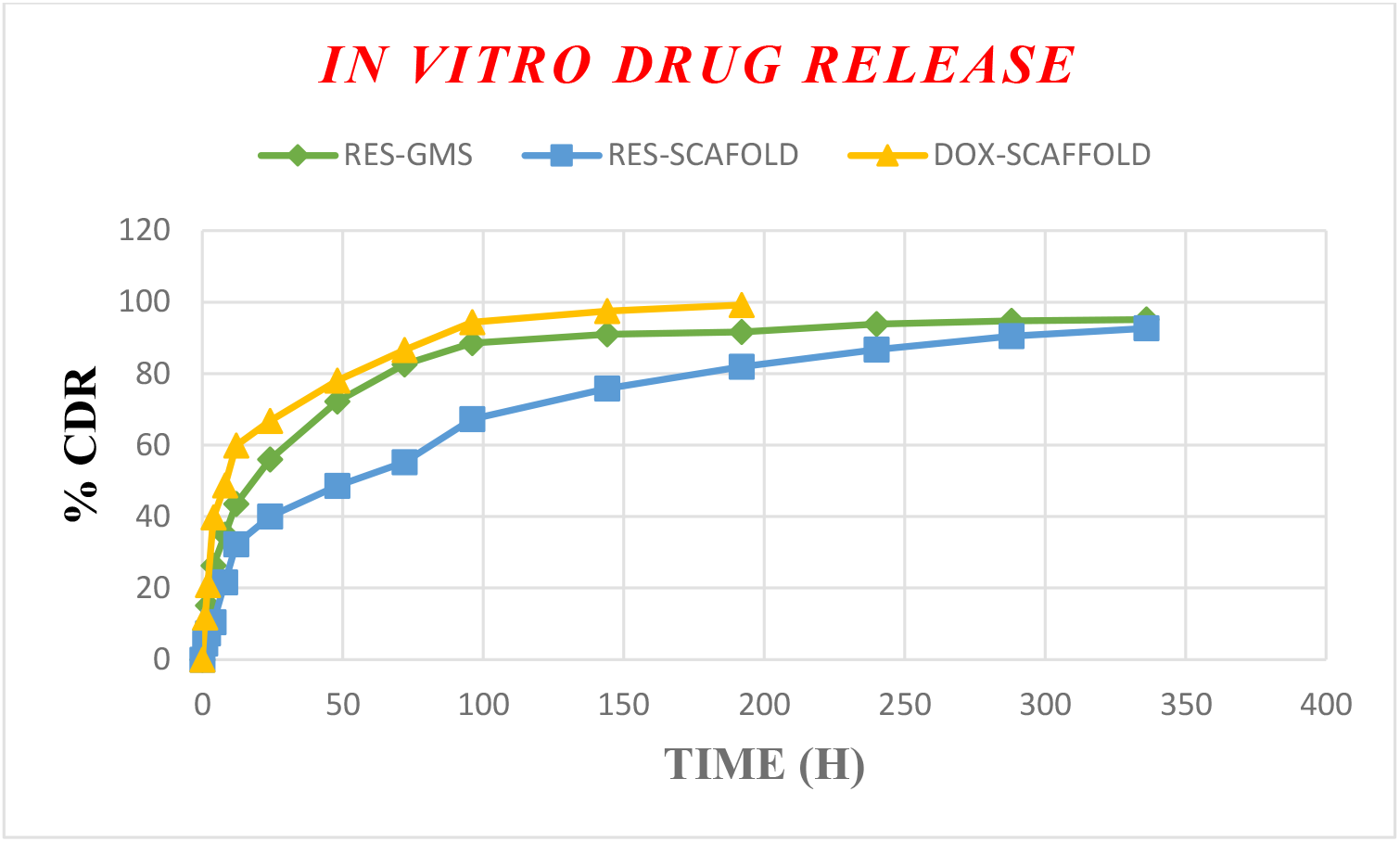
*In vitro* drug release profile of RES from RES-GMS and RES-GMS impregnated

### *In vitro* drug release Kinetics

The data acquired from in vitro drug release investigations was fitted into several release kinetic models, including zero order, first order, Higuchi, Hixson Crowell, and Korsmeyer Peppas. The Peppas model revealed that the drug release from RES-GMS had higher r^2^ values, indicating that the release of RES from the GMS was due to drug diffusion and polymer degradation. The drug release from RES-GMS-CS-CLG scaffolds, on the other hand, was shown to have higher r^2^ values for the first order and Higuchi models, indicating that drug release was concentration-dependent (control release). DOX release from the CS-CLG scaffold followed a first-order model, telling that the drug was dose-dependent **Table 2**.

**Table 2.** Release kinetics of scaffolds and microparticles

| RELEASE KINETICS |  |  |  |  |  |  |
| --- | --- | --- | --- | --- | --- | --- |
|  |  | ZERO | HIGUCHI | PEPPAS | FIRST | Hixson Crowell |
|  |  | 1 | 2 | 3 | 4 | 5 |
| RES-GMS | Slope $r^2$ | 0.430 | 5.325 | 0.256 | -0.102 | 0.160 |
|  |  | 0.626 | 0.895 | 0.9950 | 0.931 | 0.907 |
| RES-Nanohybrid Scaffold | Slope | 0.780 | 7.365 | 0.552 | -0.005 | 0.132 |
| | $r^2$ | 0.8953 | 0.985 | 0.957 | 0.990 | 0.9702 |
| DOX-Nanohybrid Scaffold | Slope | 0.350 | 7.256 | 0.314 | -0.012 | 0.016 |
| | $r^2$ | 0.818 | 0.9042 | 0.957 | 0.981 | 0.962 |

### *In vitro* Characterization

#### Antibacterial studies

It has been testified that DOX shows broad-spectral antibacterial ability and could stifle proliferation of both gram-positive and negative bacterial. The antibacterial ability mechanism by DOX is thought to be the interfering of protein synthesis inside the bacterial cells. Benefited from this advantage of DOX, DOX-loaded scaffolds also exposed the inhibition capacity of bacterial proliferation. After 24 h of incubation of scaffolds *E. coli, P.aeruginosa, MRSA* and *S. aureus*, the antibacterial feature was assessed by zone of inhibition method. Findings confirmed the excellent *in vitro* antibacterial property of DOX-loaded scaffolds against the two gram-positive and gram-negative bacterial **(Table 3)**. Increased oxidative stress is also a factor regarded as a significant cause for infection and chronic wound formation. These results showed that RES-DOX-CS-CLG scaffold illustrates excellent free radical scavenging as well as antibacterial property and might be used as an excellent wound dressing to promote wound healing.

**Table 3.** Comparative zone of inhibitions of composite scaffolds against (a) *S. aureus* (b) *MRSA* (c) *P. aeruginosa* (d) *E. coli*.

| Sl. No. | Name of the bacilli | Diameter of zone of inhibition (cms) |  |  |  |
| --- | --- | --- | --- | --- | --- |
|  |  | Placebo Scaffold | RES-GMPs Scaffold | Dox Scaffold | RES-GMS-CS-CLG scaffold |
| 1. | *E.Coli* | 0.8±0.2 | 2.2±0.2 | 3.59±0.17 | 4.20±0.3 |
| 2. | *MRSA* | 0.3±0.1 | 2.1±0.1 | 4.36±0.37 | 4.76±0.53 |
| 3. | *P.aeruginosa* | 0.2±0.1 | 2.6±0.3 | 3.78±0.15 | 4.45±0.47 |
| 4. | *S.aureus* | 0.4±0.1 | 3.2±0.2 | 3.79±0.24 | 4.31±0.36 |

#### Cytotoxicity studies (Fibroblast-Balb/3T3 cells, SRB Assay)

In vitro cytotoxicity testing of the RES-GMS-CS-CLG scaffold revealed that adding the test material to the growing medium had no effect on the viability of mouse BALB/3T3 fibroblasts. In the RES-GMS-CS-CLG scaffold construction, the fibroblasts proliferated evenly in the pores. The RES-GMS-CS-CLG scaffold’s superior biocompatibility, swelling ability, porous nature, and equal distribution of CLG, which cells have a strong affinity for, might explain this discovery. The findings indicate that the cells might grow when in contact with the scaffold and preserve their fibroblast form. This shows that the RES-GMS-CS-CLG scaffolds created are biocompatible **(Fig.8)**.

**Fig.8.**
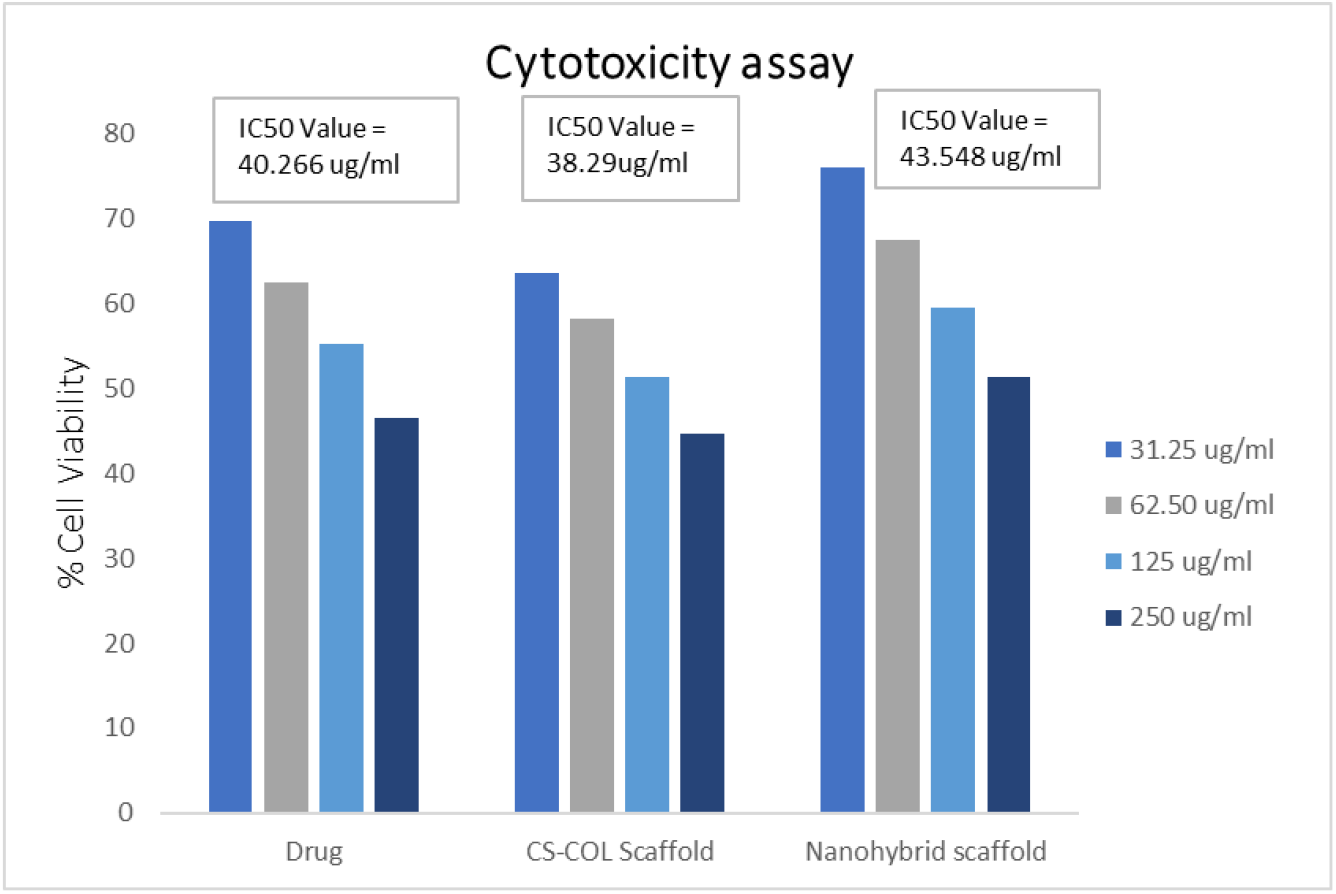
% cell viability of Balb/3T3 fibroblasts treated with drug and scaffolds

The ability of drug RES and RES-DOX-CS-CLG scaffold to promote *in vitro* wound closure was used to investigate the effects of migratory capability of Balb/3T3 fibroblast cells. Over the course of 24 hours, the rate of wound closure was monitored. Cells treated with scaffold migrated quicker than cells treated with drug and control **(Fig.9)**.

**Fig.9.**
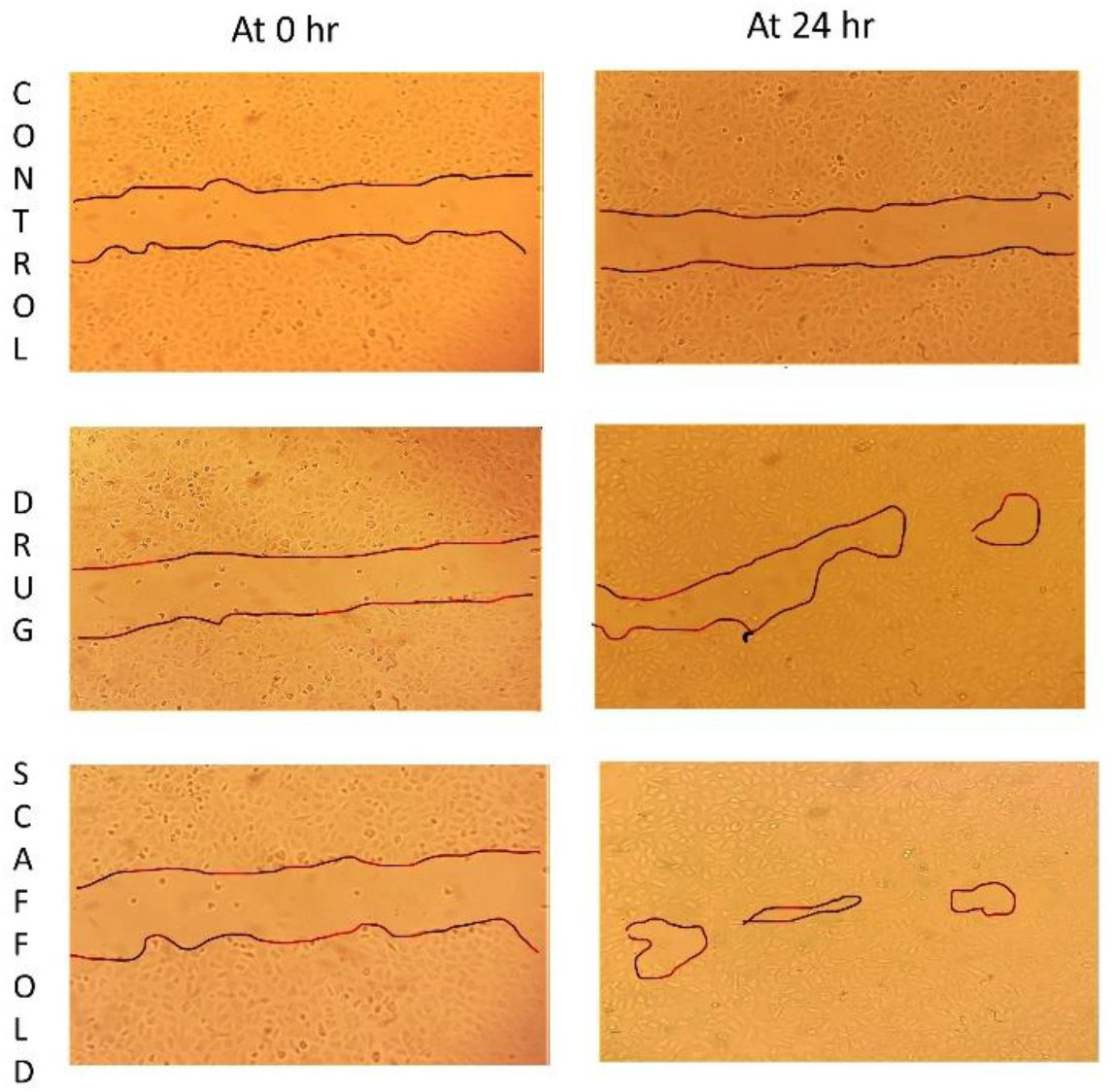
Migration of cells in Balb/3T3 fibroblasts cultured in the presence of the control, drug and scaffold

## Summary

A synergistic approach to DWH was developed in this work by combining RES-GMS and scaffold formulations, which provide a successful therapy for wound healing. SEM was used to examine RES-GMS and RES-DOX-CS-CLG scaffold. The prepared microspheres had an average particle size of 2 um and polydispersity of 0.2 to 2.9. The RES-GMS have been found to be spherical in shape with uniform size, homogeneity, and micron size The designed RES-DOX-CS-CLG scaffold has a 3D porous structure with sufficient large pore size (100-2000 m), a low degradation rate, high absorption capacity (750 percent), good mechanical strength, and the desired controlled release profile with 92% at 336h. Cell viability of Balb/3T3 cells treated with RES and RES-DOX-CS-CLG scaffold was evaluated to assess their cytotoxicity. The findings suggest that neither of them are toxic, designating that they are safe for skin wound therapy. Formulated tissue scaffolds may be valuable and viable options for wound therapy based on their *in vitro* features. More research is now being done on these scaffolds to determine the *in vivo* efficacy in appropriate animal models.

## Acknowledgement

The authors would like to thank the Department of Science and Technology Fund for Improvement of Science and Technology Infrastructure in Universities and Higher Educational Institutions (DST-FIST), New Delhi for their infrastructure support to our department.

## Ethical approval

This article does not contain any studies with human or animal subjects performed by any of the authors.

